# Tunability of DNA polymerase stability during eukaryotic DNA replication

**DOI:** 10.1101/602086

**Authors:** Jacob S. Lewis, Lisanne M. Spenkelink, Grant D. Schauer, Olga Yurieva, Stefan H. Mueller, Varsha Natarajan, Gurleen Kaur, Claire Maher, Callum Kay, Michael E. O’Donnell, Antoine M. van Oijen

**Affiliations:** Molecular Horizons and School of Chemistry and Molecular Bioscience, University of Wollongong, Wollongong, New South Wales, 2522, Australia; Illawarra Health & Medical Research Institute, Wollongong, New South Wales, 2522, Australia; Laboratory of DNA Replication, Rockefeller University, New York, NY 10065; Howard Hughes Medical Institute

**Keywords:** DNA replication, DNA, replication fork, polymerase, replisome, single molecule, fluorescence, dynamics, stability, multi-protein complexes

## Abstract

Structural and biochemical studies have revealed the basic principles of how the replisome duplicates genomic DNA, but little is known about its dynamics during DNA replication. We reconstitute the 34 proteins needed to form the *S. cerevisiae* replisome and show how changing local concentrations of the key DNA polymerases tunes the ability of the complex to efficiently recycle these proteins or to dynamically exchange them. Particularly, we demonstrate redundancy of the Pol α DNA polymerase activity in replication and show that Pol α primase and the lagging-strand Pol δ can be re-used within the replisome to support the synthesis of large numbers of Okazaki fragments. This unexpected malleability of the replisome might allow it to deal with barriers and resource challenges during replication of large genomes.

## Introduction

To robustly synthesize genomic DNA, the eukaryotic replisome requires a large number of interacting protein factors with different enzymatic activities. Key components include the 11-subunit CMG helicase and three different multi-subunit B-family DNA polymerases – the leading-strand Pol ε, lagging-strand Pol δ, and Pol α-primase (Bell and Labib, 2016; Burgers and Kunkel, 2017). CMG unwinds DNA by translocating along one of the strands in a 3’ to 5’ direction while forming a complex with Pol ε (Langston et al., 2014; Sun et al., 2015) to support highly processive synthesis of DNA on the leading strand. On the lagging strand, Pol α generates ~25-nt RNA–DNA primers that Pol δ extends to generate ~150-bp Okazaki fragments (Bell and Labib, 2016). The commonly accepted model of replication depicts Pol ε stably anchored to CMG but shows Pol δ not physically tethered to the replisome and subsequently replaced for the synthesis of each Okazaki fragment (Bell and Labib, 2016). Biochemical and structural studies have provided valuable insights into basic enzymatic activities and overall architecture of the eukaryotic replisome. However, the dynamic behavior of the various replisomal components is largely unexplored due to the challenges associated with the averaging over ensembles of molecules that is needed to gain structural and functional insight.

We demonstrate here the reconstitution of the *S. cerevisiae* replisome by purified protein factors and the visualization of processive DNA replication at the single-molecule level. We show that all three replicative polymerases can remain stably associated with the replisome for the synthesis of tens of kilobases (kb), and that the DNA synthesis activity of Pol α is dispensable under these conditions. Pol δ is retained at the fork while synthesizing large numbers of successive Okazaki fragments. This unexpected observation of recycling of the lagging-strand Pol δ without dissocation from the replisome directly challenges textbook models and implies physical connectivity between Pol δ and the replisome, and the formation of loops in the lagging-strand DNA. We demonstrate that this retention of Pol δ is facilitated through interaction between the Pol 32 subunit of Pol δ and the Pol 1 subunit of Pol α (Huang et al., 1999; Johansson et al., 2004). Interestingly, when challenged with competing polymerases, both Pol ε and Pol δ are able to exchange from solution into a moving replisome in a concentration-dependent manner. We propose this balance between replisome stability and plasticity enables the complex to provide access to other binding partners during S phase while not compromising the stability and robustness of the replisome.

## Results

### Single-molecule DNA replication

We visualized individual budding yeast replisomes using a real-time single-molecule fluorescence assay that allows us to monitor simultaneously DNA synthesis and protein dynamics. We assemble linear, double-stranded DNA (dsDNA) molecules (18.3 kb length) in a microfluidic flow cell placed onto a fluorescence microscope. The DNA is stretched and attached to the surface at both ends (Figure S1A,B). A pre-made synthetic replication fork at one end of the DNA enables direct loading of the replisomal proteins onto the DNA (Figure 1A, left). DNA synthesis is initiated by loading CMG and the Mcm10 initiation factor (CMGM) onto the template followed by the introduction of Ctf4, Mcm10, Mrc1–Tof1–Csm3 (MTC), PCNA, RFC, RPA, DNA polymerases α, δ, ε, Mg^2+^, the four dNTPs and four rNTPs (Figure 1A, right) (Georgescu et al., 2015). Real-time synthesis trajectories were obtained from near-TIRF fluorescence imaging of Sytox-Orange (S.O.) stained dsDNA in the absence of buffer flow. As DNA synthesis proceeds the leading strand appears as a diffraction-limited spot that moves unidirectionally along the template DNA. Initially weak in intensity, the spot increases in intensity as more dsDNA is generated at the leading strand and accumulates into a collapsed globular coil with dimensions smaller than the diffraction-limited resolution of the imaging system (Figure 1B, left; Video S1). The intensity per base pair of this coiled leading-strand product is similar to the intensity per base pair as measured over the stretched template (Figure S1C,D). To establish that the observed events correspond to simultaneous leading- and lagging-strand synthesis, we divided the DNA template into three regions — the leading-strand spot (‘Lead’), the length of DNA behind it (‘Lag’) and the length ahead of it (‘Parental’) (Figure 1B, right). For every time point, the DNA content is calculated from the integrated fluorescence intensity in each region (Ganji et al., 2018). During replication, the DNA content of the parental region decreased while DNA content of the leading- and lagging-strand regions increased simultaneously (Figure 1B right; Figure S1E). Importantly, in the absence of either Mg^2+^, nucleotides, CMG, or DNA polymerases replication events were not observed (Figure S2A).

**Figure 1.**
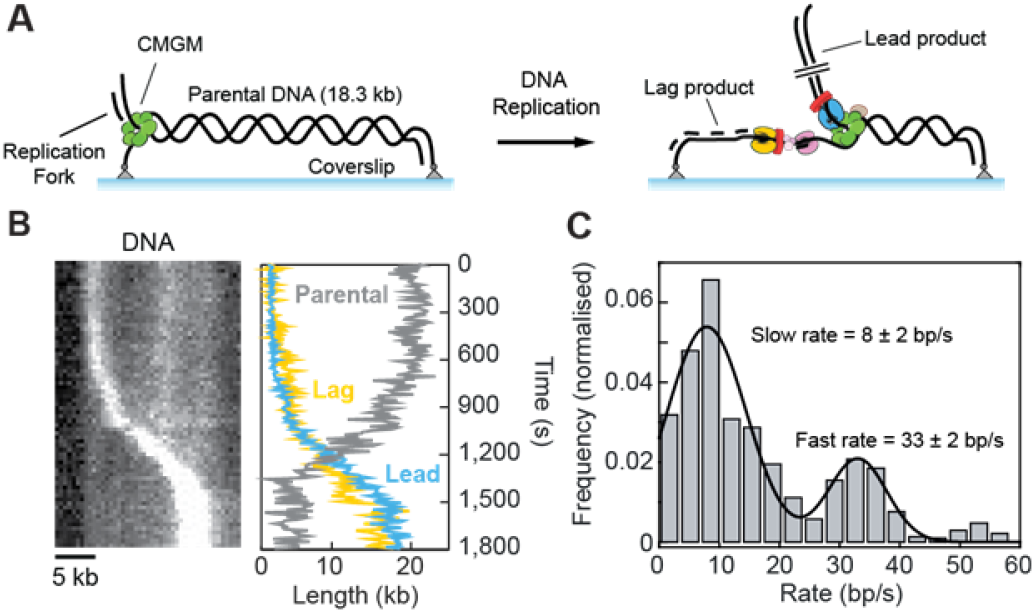
Single-molecule visualization of DNA synthesis. (A) Schematic representation of DNA-replication assay (see methods for details). (B) (left) Kymograph showing DNA replication on a single DNA substrate. The leading-strand tail appears as a bright spot that moves in a unidirectional manner, while simultaneously increasing in intensity. (right) Length of the lagging strand (‘Lag’), leading strand (‘Lead’) and parental DNA (‘Parental’) as a function of time, measured by the integrated intensity of the dsDNA. (C) Single-molecule rate distribution. The bimodal distribution was fit with the sum of two Gaussians (black line) with rates of 8 ± 2 bp/s and 33 ± 2 bp/s (*n* = 96 molecules). Errors represent S.E.M.

To quantify the instantaneous rates of replication, we tracked the position of the leading-strand spot (Tinevez et al., 2017). The measured population-averaged rate of 19 ± 6 bp/s (mean ± S.E.M.; *n* = 96 molecules) (Figure S2B,C) is consistent with previously reported ensemble *in vitro* and *in vivo* measurements (Aria and Yeeles, 2018; Hodgson et al., 2007; Sekedat et al., 2010; Szyjka et al., 2005; Tourriere et al., 2005; Yeeles et al., 2017). Rates of DNA synthesis varied within individual replisomes (Figure 1C, Figure S2B,C), with the single-molecule rate distribution having two distinct peaks at 8 ± 2 bp/s and 33 ± 2 bp/s (mean ± S.E.M.; *n* = 96 molecules, 315 segments). This bimodal rate distribution was reported in our previous single-molecule studies of leading-strand synthesis and shown to correspond to dynamic interaction of the MTC complex with the replisome (Lewis et al., 2017b).

### Pre-assembled replisomes replicate DNA as a highly stable entity

To measure the stability of synthesizing replisomes on DNA, we carried out single-molecule pre-assembly replication assays. In these assays, the replisome is first pre-assembled on surface tethered DNA in the flow cell (Figure 2A, left and middle). Next, the flow cell is washed and replication is initiated by introduction of a replication solution that omits all three polymerases. This protocol ensures only the initially assembled polymerases remain, eliminating the possibility of other polymerases associating with the replisome (Figure 2A, right). We hypothesized that the requirement for a new Pol δ for each Okazaki fragment would prevent processive synthesis. Surprisingly, these conditions support processive DNA replication, with synthesis rates similar to those measured with excess polymerases in solution (Figure 2B; Figure S2C). This observation suggests that Pol δ can be stably associated to the replisome, challenging the current view that a new Pol δ holoenzyme is recruited to extend each Okazaki fragment (Bell and Labib, 2016). These results also reveal that the Pol α-primase is stably associated to the replication machinery and primes multiple Okazaki fragments.

**Figure 2.**
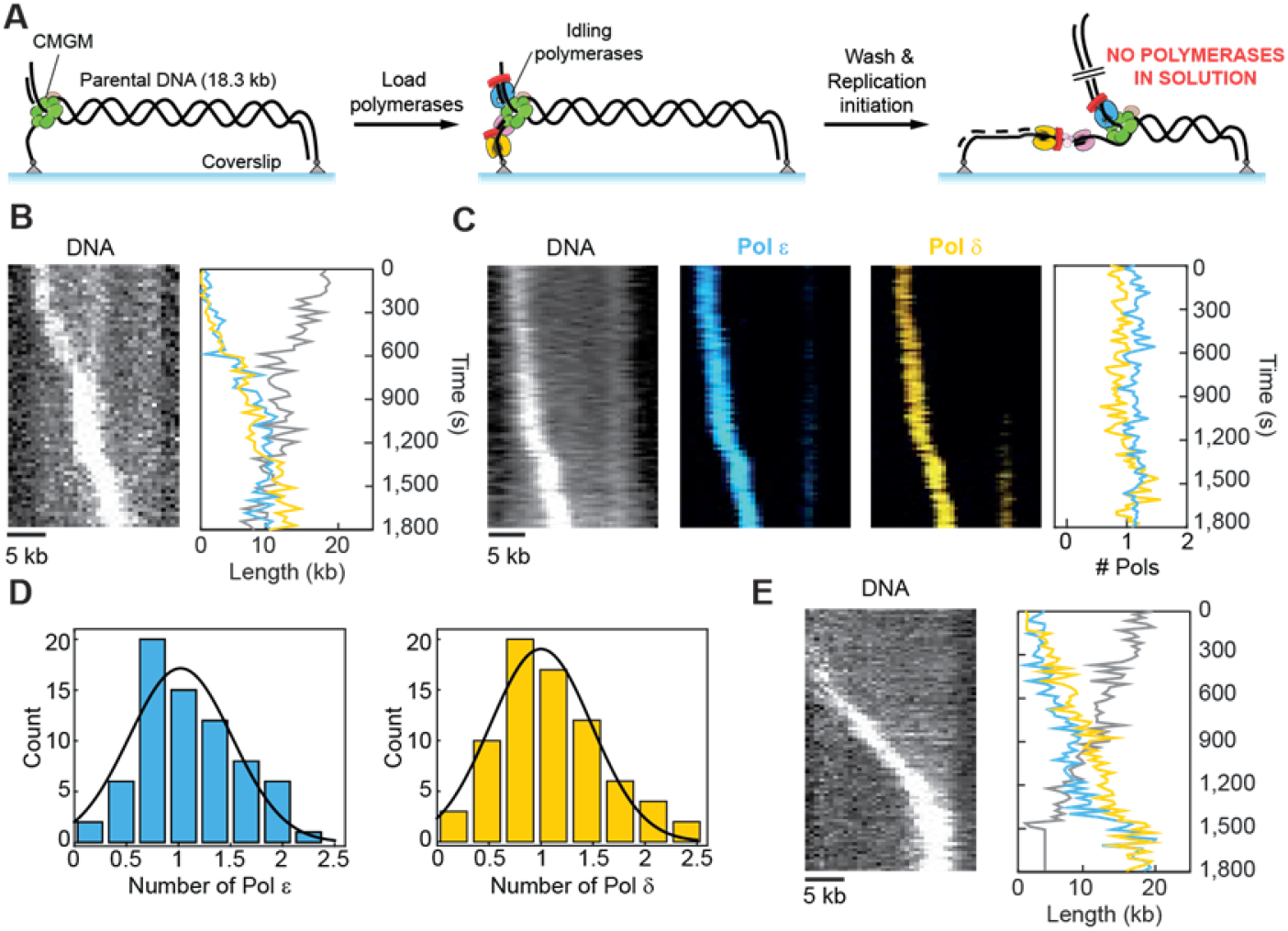
Direct visualization of polymerase stability during processive DNA synthesis. (A) Schematic representation of the pre-assembly DNA replication assay (see methods for details). (B) (left) Kymograph showing activity of a pre-assembled replisome on a single DNA substrate in the absence of polymerases in solution. (right) Length of the lagging strand (yellow), leading strand (blue) and parental DNA (gray) as a function of time. (C) Example kymographs showing the DNA, labeled Pol ε, labeled Pol δ, and the polymerase intensities as a function of time. Both polymerases co-localize with the leading-strand spot. (D) Distribution of the number of Pol ε (blue) and Pol δ (yellow) at the fork (*n* = 70). A Gaussian fit (black line) gives 1.0 ± 0.2 (mean ± S.E.M.) and 1.0 ± 0.2 (mean ± S.E.M.) respectively. (E) (left) Kymograph showing DNA replication of pre-assembled replisomes containing Pol α^Cat^. (right) Length of the lagging strand (yellow), leading strand (blue) and parental DNA (gray) as a function of time.

### Direct visualization confirms presence of a single Pol δ in a processive replisome

To further explore the observation that Pol δ remains tethered to the replisome, we repeated our single-molecule pre-assembly assay in the presence of fluorescently labeled Pol ε and Pol δ (Figure 2C). Labeling did not affect the average rates of DNA synthesis (Figure S2C). The kymographs in Figure 2C show bright fluorescent spots for both the labeled Pol ε and Pol δ during DNA synthesis. Both DNA polymerases also co-localize with the leading-strand spot, consistent with both polymerases stably incorporated into reconstituted replisomes. To determine the stoichiometry of Pol ε and Pol δ, we divided the intensity at the fork by the intensity of a single polymerase. We observe the continuous presence of one Pol ε and one Pol δ at actively synthesising replication forks (Figure 2D). DNA synthesis on the lagging strand is supported by both Pol δ and the DNA polymerase activity of the multifunctional Pol α. To investigate the influence of DNA synthesis activity by Pol α on the lagging strand, we repeated the pre-assembly assay with a mutant of Pol α-primase that is able to produce RNA primers but is unable to extend these into DNA (Pol α^Cat^, Fig S3). Remarkably, the DNA polymerase activity of Pol α is dispensable during processive leading- and lagging strand synthesis (Figure 2E, Figure S2C). Taken together, these observations are consistent with a model in which Pol δ supports lagging-strand synthesis (Aria and Yeeles, 2018; Yeeles et al., 2017) and Pol α is only required for its primase activity.

### Direct visualization of concentration-dependent exchange of Pol ε

Our observations and previous studies (Douglas et al., 2018; Langston et al., 2014; Lewis et al., 2017b) suggest that Pol ε is a stable component of the replisome. However, recent single-molecule experiments have demonstrated that stable components of multi-protein complexes can undergo dynamic exchange when challenged with competing binding partners (Delalez et al.; Graham et al.; Li et al.; Scherr et al.). In particular, complexes held together by multiple weak interactions support high stability in the absence of competing factors, but rapid exchange in the presence thereof (van Oijen and Dixon, 2015). We hypothesized that the multiple contact points between Pol ε and the replisome may similarly allow it to exchange dynamically in the presence of free polymerases in solution (Aberg et al., 2016). To visualize such polymerase dynamics we repeated the assay but now with labeled polymerases in solution (Figure 3A). The presence of excess polymerases in solution did not result in changes to replication kinetics (Figure S2C). In the presence of all three polymerases in solution, we detect on average one Pol ε and one Pol δ present at the replication fork (Figure 3B). Interestingly, in 86 ± 5% (mean ± S.E.M.; *n* = 55) of events DNA synthesis begins only after binding of Pol δ, consistent with a model where physical tethering between the replisome, Pol α, and Pol δ allows processive Okazaki fragment synthesis to occur only after recruitment of all required polymerases.

**Figure 3.**
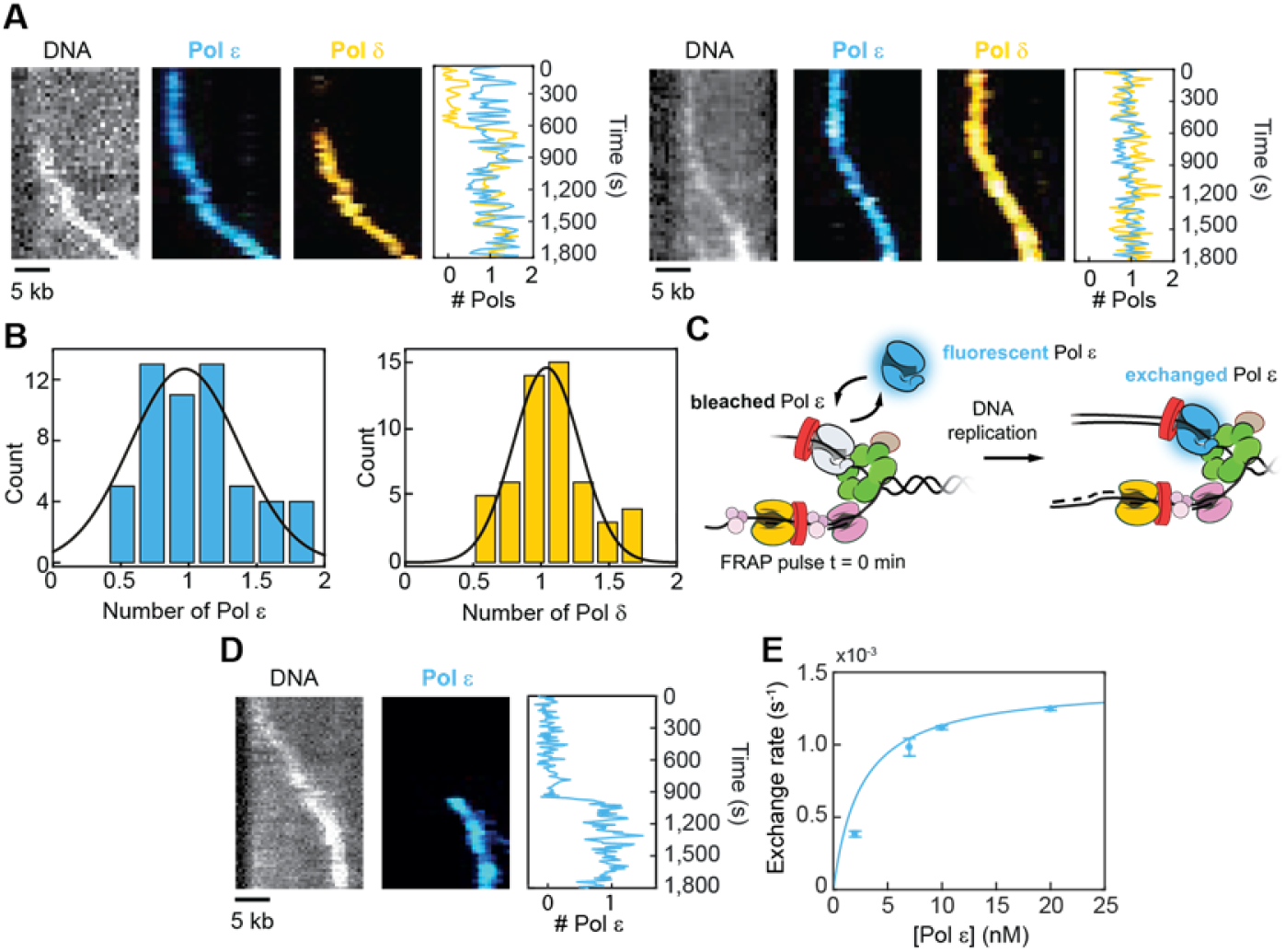
Visualization of Pol ε dynamics. (A) Example kymographs showing the DNA, labeled Pol ε, labeled Pol δ, and the polymerase intensities as a function of time. Both polymerases co-localize with the leading-strand spot. (B) Distributions of the number of Pol ε (blue) and Pol δ (yellow) at the fork (*n* = 55). A Gaussian fit (black line) gives 1.0 ± 0.2 (mean ± S.E.M.) and 1.0 ± 0.1 (mean ± S.E.M.) respectively. (C) Schematic representation of the FRAP assay. (D) Example kymograph showing exchange of Pol ε at the fork with DNA (left), labeled Pol ε (middle), and the labeled Pol ε intensity. (E) Exchange rate as a function of Pol ε concentration. The line represents a hyperbolic fit, giving a maximum exchange rate of (1.4 ± 0.5) ×10^−3^ s^−1^ (mean ± error of the fit).

Next, to identify whether exchange occurs, we carried out single-molecule FRAP (Fluorescence Recovery After Photobleaching) experiments (Beattie et al., 2017; Lewis et al., 2017a; Spenkelink et al., 2019). Using labeled Pol ε, we photobleached all labeled polymerases at the replication fork using a pulse of high laser intensity after initiation of replication (Figure 3C, left). After bleaching, we monitored fluorescence recovery to see if new unbleached Pol ε from solution could exchange into the replisome (Figure 3C, right). Figure 3D shows a kymograph of the fluorescence recovery of Pol ε in the presence of 20 nM Pol ε in solution. We obtained an average exchange rate of ~1 per 10 min for these conditions (i.e. (12.5 ± 0.1) ×10^−4^ s^−1^, Figure S4A). This rate is equivalent to exchange of one Pol ε over a length of ~15 kb. Decreasing the labeled Pol ε concentration from 20 to 2 nM, the mean exchange rate decreased (Figure 3E). These results demonstrate that Pol ε undergoes concentration-dependent exchange at the replication fork.

As an internal control, we set out to characterize the exchange kinetics of the CMG helicase (Figure 4A). A number of mechanisms are in place during the cell cycle to achieve the precise loading of exactly one CMG per replication fork (Abid Ali et al., 2017; Bell and Labib, 2016; Douglas et al., 2018). Combined with structural data that point to a role of CMG as the central organizer of the replisome, we hypothesized CMG to remain stably bound during elongation. Using fluorescently labeled CMG, we observe a single CMG remaining stably bound during DNA synthesis on time scales longer than 30 min, even when challenged with excess CMG in solution (Figure 4B and C).

**Figure 4.**
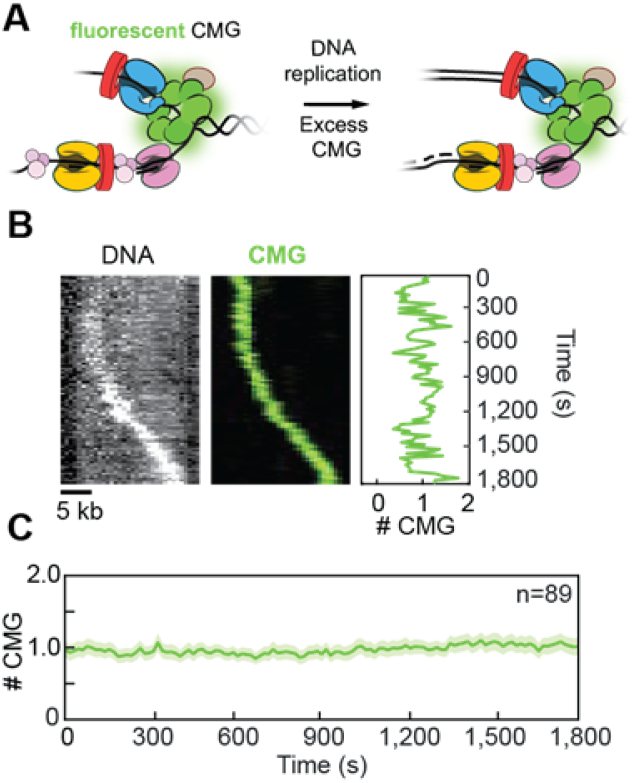
CMG is a stable component during replication. (A) Schematic representation of the assay. (B) Example kymograph showing the DNA, labeled CMG, and the CMG intensity as a function of time. CMG co-localizes with the leading-strand DNA spot. (C). Average number of CMG molecules at elongating replication forks over time, giving 1.0 ± 0.7 (mean ± S.E.M).

### Direct visualization of Pol δ dynamics

To test whether Pol δ displays exchange behavior similar to Pol ε, we repeated the FRAP experiments with labeled Pol δ (Figure 5A, Figure S4B). We observe the exchange rate of Pol δ is also dependent on the concentration of Pol δ in solution (Figure 5E). Importantly, even at the highest concentration of Pol δ that still allows visualization of single molecules (20 nM), the Pol δ exchange rate is such that it would correspond to the synthesis of many Okazaki fragments. To understand how the Pol δ exchange rate compares to the Okazaki-fragment cycling time, we quantified the average length of Okazaki fragments under our experimental conditions. We used fluorescently labeled RPA to assess the amount of ssDNA as a measure of the Okazaki fragment size (Figure 5B, left). As expected, RPA was always localized at the replication fork. Consistent with biochemical studies (Georgescu et al., 2015) we observe that the number of RPA molecules is dependent on the concentration of Pol α (Figure 5B, right). Given that the footprint of RPA is 30 ± 2 nt (Figure 5C) and we see 3.7 ± 0.5 (mean ± S.E.M.; *n* = 64) RPA molecules at the fork, we determine that the average Okazaki fragment length in our single-molecule assay is 111 ± 16 bp (mean ± S.E.M.), consistent with *in vivo* studies (Bell and Labib, 2016). At an average replication rate of 19 ± 6 bp/s our observation suggests that Pol δ is retained within the replisome for synthesis of 142 ± 78 Okazaki fragments, and therefore must have stabilising contacts with other replisomal components.

**Figure 5.**
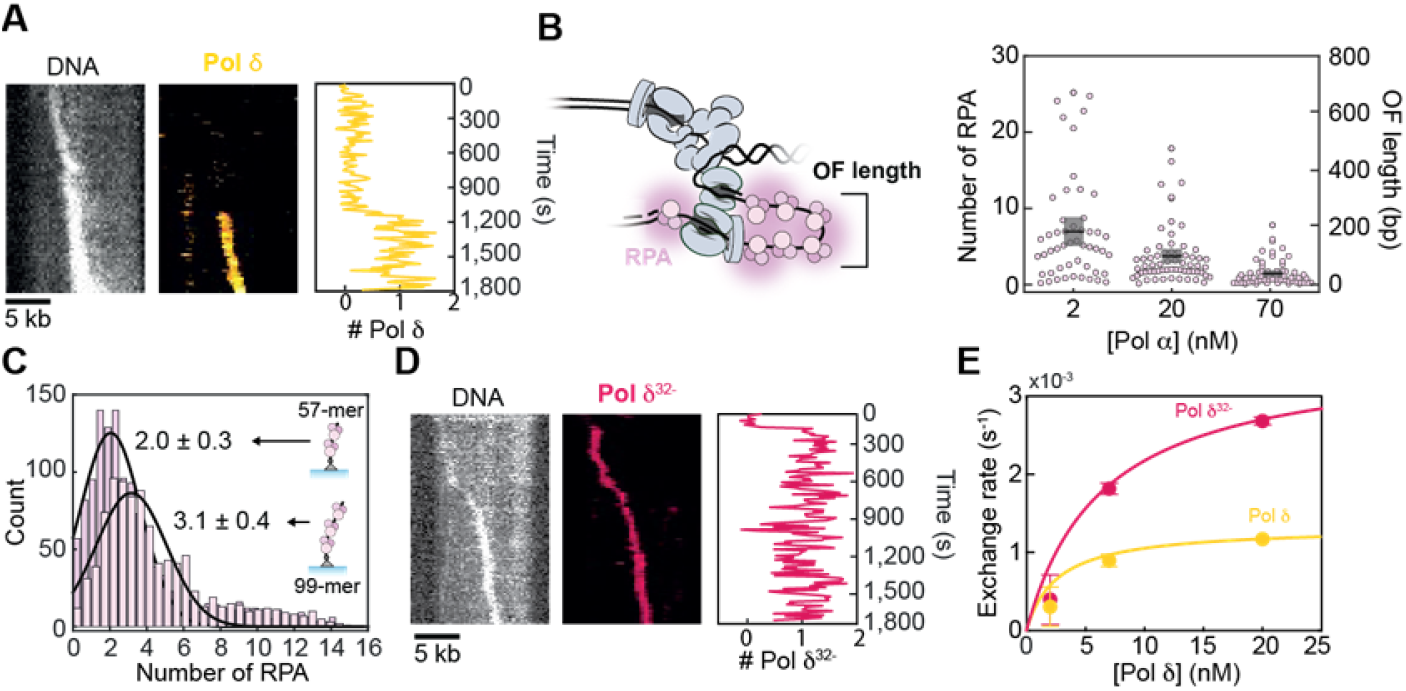
Pol δ dynamics. (A) Example kymograph showing the DNA and labeled Pol δ, and corresponding labeled Pol δ intensity at the fork. (B) (left) Schematic representation showing the relationship between the number of RPA molecules and Okazaki-fragment length. (right) Scatter plots of the number of RPA for 2 nM (7.0 ± 0.9; *n* = 51), 20 nM (3.7 ± 0.5; *n* = 64), and 70 nM (1.5 ± 0.3; *n* = 60) Pol α. The black line and gray box represent the mean and S.E.M. (C) Histograms of the number of RPA molecules binding to either a 57-mer (purple, *n* = 1299) and 99-mer oligo (pink, *n* = 939). The average ssDNA–RPA footprint is 30 ± 2 nt. Error represents S.E.M. (D) Example kymograph showing the DNA and labeled Pol δ^32-^, and corresponding labeled Pol δ^32-^ intensity at the fork. (E) Exchange rate as a function of polymerase concentration with Pol δ (yellow) and Pol δ^32-^ (magenta). The lines represent hyperbolic fits, giving a maximum exchange rate of (1 ± 3) ×10^−3^ (mean ± error of the fit) for Pol δ (yellow) and (4 ± 8)×10^−3^ (mean ± error of the fit) Pol δ^32-^ (magenta).

We then set out to identify the mechanism through which Pol δ is retained in replisomes. The Pol 32 subunit of Pol δ is documented to interact with the Pol 1 subunit of Pol α (Huang et al., 1999; Johansson et al., 2004). Pol α also binds to CMG through an interaction with Ctf4 and Mcm10 (Simon et al., 2014; Warren et al., 2009). Thus, we predicted that elimination of contact between Pol 32 of Pol δ and Pol α would result in a change in exchange of the Pol δ into the replisome. If the Pol 32–Pol 1 interaction is important, the exchange of Pol δ^32-^ (Pol δ lacking the Pol 32 subunit) should be measurably faster than the exchange of the complete Pol δ holoenzyme. To test this hypothesis, we fluorescently labeled Pol δ^32-^ and repeated the FRAP measurements (Figure 5D, Figure S4C). Indeed, the exchange rate of Pol δ^32-^ is ~2.5-fold faster than the exchange of Pol δ holoenzyme ((27.0 ± 0.5) ×10^−4^ s^−1^ compared to (12 ± 5) ×10^−4^ s^−1^) (Figure 5E). These data show that Pol α helps retain Pol δ in the replisome.

## Discussion

We report here the reconstitution and visualization at the single-molecule level of DNA synthesis by budding yeast replisomes. Our real-time fluorescence assay allows us to directly visualize replication kinetics and quantify protein dynamics in individual replisomes – observables that are not accessible *via* classical biochemical approaches. The average observed replication rates are similar to those previously reported in ensemble biochemical reactions (Aria and Yeeles, 2018; Georgescu et al., 2015; Kurat et al., 2017; Yeeles et al., 2017) and are within the range of replication fork rates inside the cell (Conti et al., 2007; Hodgson et al., 2007; Sekedat et al., 2010). We show that the eukaryotic replisome acts as a stable processive machine under dilute conditions. In the presence of excess polymerases, however, the replisome is not fixed in composition, but instead is a highly dynamic complex continually exchanging major components. This model is founded upon three observations. (i) In the absence of all three replicative polymerases in solution the replisome forms a stable complex able to support processive, concerted leading- and lagging-strand synthesis (Figure 2). (ii) Pol δ can be retained at the replication fork for multiple Okazaki fragments, mediated at least in part through an interaction with Pol α (Figure 4). (iii) The leading- and lagging-strand polymerases Pol ε and Pol δ exchange during DNA synthesis in a concentrationdependent manner without affecting replication rate (Figure 3 and 4, Figure S2C).

In contrast to long-standing views that replisome architecture is static, single-molecule fluorescence experiments have documented dynamic exchange of components at physiologically relevant time scales in large multi-protein complexes across all domains of life (Geertsema et al., 2014; Liao et al.). Our work provides the first direct evidence that Pols ε and δ are similarly exchanged from solution in a concentration-dependent manner during DNA synthesis. In the absence of polymerases in solution, the original polymerases are retained and the replisome forms a highly stable complex resistant to dilution (Figure 6A). In the presence of excess polymerases, they are exchanged into the replisome at a rate dependent on their concentration (Figure 6B). Concentration-dependent exchange can be rationalized through an interaction network consisting of multiple weak interactions. Under dilute conditions, transient disruption of any one of these interactions would be followed by its rapid reformation to prevent dissociation. If, however, there are exogenous competitors in close proximity to the complex, one of these can bind at a transiently vacated binding site and consequently be at a sufficiently high local concentration to compete out the original protein.

**Figure 6.**
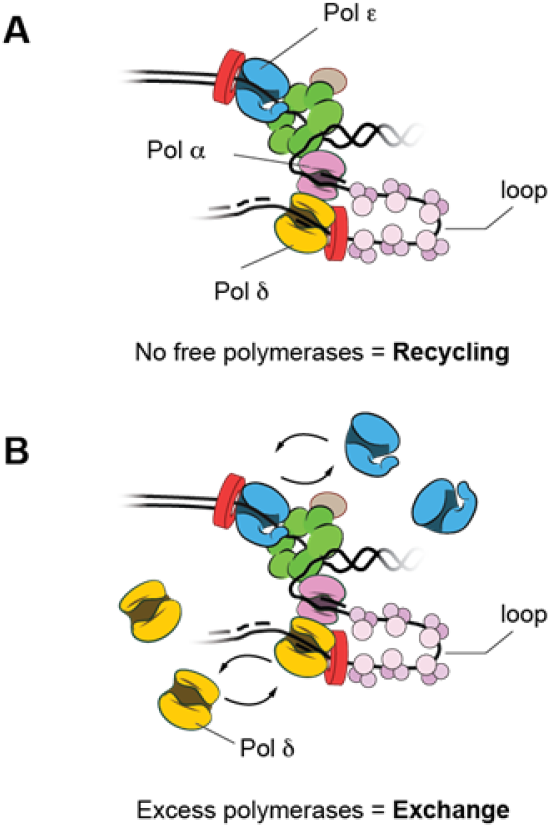
Compositional plasticity at the eukaryotic replication fork. (A) All three replicative polymerases can be recycled forming a stable complex in the absence of free polymerases in solution. Pol δ is linked to CMG through interaction with Pol α through its Pol 32 subunit. (B) Pol ε and Pol δ exchange in a concentration-dependent manner.

A concentration-dependent exchange mechanism likely plays an important role in genomic integrity. Through such a mechanism, the replisome has access to a plurality of molecular pathways to achieve and ensure continuous replisome progression under a variety of cellular stresses. Re-priming of leading-strand synthesis past DNA damage is highly inefficient (Taylor and Yeeles, 2018). As a result, the replisome will need to recruit specialized repair polymerases directly to the fork for efficient bypass of damage. Concentration-dependent exchange of Pol ε on the leading strand provides a simple molecular mechanism to grant repair polymerases access to the replisome during the initial response to DNA damage.

Dynamic exchange of polymerases also provides the replisome with a straightforward way to pass through cohesion rings that hold together replicated sister chromatids. Cohesin can move over obstacles ~11 nm in size (Stigler et al., 2016). Since the size of the replisome is much larger (Sun et al., 2015) it is difficult to envisage how it can pass through cohesin rings. We hypothesize that CMG (~10 nm) may fit through the pore and that dynamic exchange of other components allows the replisome to pass as a minimal complex to be rejoined with its polymerases immediately after.

It is generally assumed that Pol δ is not associated with CMG and that a new Pol δ is recruited for the synthesis of each successive Okazaki fragment. In contrast, our results show that Pol δ can be retained at the fork for synthesis of multiple Okazaki fragments. This observation implies that there are one or more interactions between Pol δ and a stable part of the replisome. We discover that one of these interactions is with Pol α through the Pol 32 domain of Pol δ — without this domain, Pol δ exchange is ~2.5-fold faster (Figure 5). Pol α has a specific interaction with CMG (Georgescu et al., 2015) and can be tethered to CMG *via* interactions with Ctf4 and Mcm10 (Gambus et al., 2009; Warren et al., 2009). We, therefore, propose that Pol δ is tethered to CMG mediated by the interaction with Pol α. Retention of Pol δ over the time scales presented here, combined with the RPA stoichiometry, implies that lagging-strand replication loops can be formed at the eukaryotic replication fork (Figure 6) (Chastain et al., 2003). Additionally, the redundancy of the DNA polymerase acitvity of Pol α, suggests it may perform other functions in the cell other than priming in DNA replication.

## Acknowledgements

The authors thank Dan Zhang for purification of CMG, Cees Dekker for pSuperCos1, Daniel Zalami for his help in setting up Trackmate, and Karl Duderstadt for critical reading of the manuscript. This work was supported by Australian Research Council Grant DP180100858, Australian Laureate Fellowship FL140100027 (to A.M.v.O.), NIH Grant GM-115809 (to M.E.O.), Howard Hughes Medical Institute (to M.E.O.), T32 CA009673 (to G.S.) and K99 GM126143 (to G.S.).

## Author contributions

Conceptualization, J.S.L., L.M.S., G.D.S., M.E.O., A.M.v.O.; Formal analysis, J.S.L., L.M.S., S.H.M.; Funding acquisition, M.E.O., A.M.v.O.; Investigation, J.S.L., L.M.S., G.D.S., S.H.M., V.N., C.M., C.K.; Methodology, J.S.L., L.M.S., A.M.v.O.; Resources, G.D.S., O.Y., G.K.; Software, L.M.S.; Supervision, M.E.O., A.M.v.O.; Validation, J.S.L., L.M.S.; Visualization, J.S.L., L.M.S., A.M.v.O.; Writing – original draft, J.S.L., L.M.S., G.D.S., M.E.O., A.M.v.O.

## STAR METHODS

### KEY RESOURCES TABLE

| REAGENT or RESOURCE | SOURCE | IDENTIFIER |
| --- | --- | --- |
| Bacterial and Virus Strains |  |  |
| *S. cerevisiae* strain W303 | Brian Chait Lab<br>(Rockefeller University) | N/A |
| *Escherichia coli* BL21-DE3 codon plus RIL | Agilent | Cat #230245 |
| Chemicals, Peptides, and Recombinant Proteins |  |  |
| Alexa Fluor 647 NHS ester mono-reactive dye | Life Technologies | Cat# A20006 |
| Alexa Fluor 488 maleimide mono-reactive dye | Life Technologies | Cat# A10254 |
| Cy5 NHS ester mono-reactive dye | GE Healthcare | Cat# PA15101 |
| LD650 maleimide mono-reactive dye | Lumidyne Technologies | Cat# LD650-MAL |
| Coenzyme A trilithium salt | Sigma-Aldrich | Cat# C3019 |
| d-desthiobiotin | Sigma-Aldrich | Cat# D1411 |
| Chloroquine diphosphate salt | Sigma-Aldrich | Cat# C6628 |
| SYBR-Safe DNA stain | Life Technologies | Cat# S33102 |
| SYTOX Orange nucleic acid stain | Life Technologies | Cat#S11368 |
| γ-<sup>32</sup>P-ATP | Perkin Elmer | Cat# NEG002A |
| Polydimethylsiloxane (184 Silicone Elastomer) | Sylgard | Cat# 500GM184KI T |
| Biotin-PEG and MPEG (MW 5,000) bundle | Laysan Bio | Cat# BIO-PEG-SVA-5K-100MG & MPEG-SVA-5K-1g |
| 3-Aminopropyltriethoxysilane (APTES) | Alfa Aesar | Cat# A10668 |
| dNTP Bundle | Jena Bioscience | Cat# NU-1005L |
| ATP solution | Jena Bioscience | Cat# NU-1010 |
| NTP Bundle | Jena Bioscience | Cat# NU-1014L |
| 3 x Flag-tag | EZBiolab | Cat# cp7204 |
| Restriction endonuclease *Bst*XI | New England Biolabs | Cat# R0113L |
| T4 DNA ligase | New England Biolabs | Cat# M0202L |
| T4 Polynucleotide Kinase | New England Biolabs | Cat# M0201L |
| NeutrAvidin protein | Life Technologies | Cat# 31000 |
| PMSF Protease Inhibitor | ThermoFisher | Cat# 36978 |
| CMG helicase | (Langston et al., 2014) | N/A |
| Pol $\epsilon$ | (Langston et al., 2014) | N/A |
| RFC clamp loader | (Langston et al., 2014) | N/A |
| PCNA | (Langston et al., 2014) | N/A |
| RPA | (Langston et al., 2014) | N/A |
| Mcm10 | (Langston et al., 2017; Lewis et al., 2017b) | N/A |
| MTC | (Langston et al., 2017; Lewis et al., 2017b) | N/A |
| Pol $\alpha$ | (Georgescu et al., 2014) | N/A |
| Pol $\delta$ | (Georgescu et al., 2014) | N/A |
| Ctf4 | (Georgescu et al., 2014) | N/A |
| Pol $\alpha^{CAT}$ | This study | N/A |
| LD650/AF488–Pol $\epsilon$ | This study | N/A |
| LD650–CMG | (Wasserman et al., 2019) | N/A |
| Pol $\delta^{32-}$ | This study | N/A |
| Oligonucleotides |  |  |
| See Table S1 | IDT | N/A |
| Plasmid DNA |  |  |
| Sfp pet29b C-terminal His Tag | Addgene | Cat# 75015 |
| pSuperCos1 | Cees Dekker Lab (TU Delft) | N/A |
| Software and Algorithms |  |  |
| MATLAB 2016b | MathWorks | https://www.mathworks.com/products/matlab.html |
| ImageJ/FIJI (1.51w) | (Schindelin et al.) | https://imagej.net/Fiji |
| TrackMate v3.6.0 | (Tinevez et al., 2017) | N/A |

### CONTACT FOR REAGENT AND RESOURCE SHARING

Further information and request for resources and reagents should be directed to and will be fulfilled by the Lead Contact, Antoine van Oijen.

### EXPERIMENTAL MODEL AND SUBJECT DETAILS

#### Yeast and bacterial strains

Proteins were purified from *Saccharomyces cerevisiae* strain W303 (genotype: *ade2-1 ura3-1 his3-11,15 trp1-1 leu2-3,112 can1-100 bar1Δ MATa pep4::KANMX6*) or *Escherichia coli* BL21-DE3 codon plus RIL cells transformed with plasmids for overexpression of proteins of interest as detailed in the Key resources table.

### METHOD DETAILS

#### Protein purification and labeling

##### Purification of Pol δ^32-^

Pol δ was purified as previously described (Georgescu et al., 2014), the final step of which involves an elution from a sulphopropyl cation exchange column (GE Healthcare) with a 100-500 mM NaCl gradient in a buffer containing 350 mM potassium glutamate and 25 mM HEPES pH 7.5. Whereas, Pol δ (as a complete holoenzyme) elutes around 350 mM NaCl, Pol δ^32-^ elutes around 250 mM NaCl, allowing complete separation (Figure S5). Thus, peak fractions at 250 mM NaCl were aliquoted, flash frozen, and stored at −80°C.

##### Purification of Pol α^Cat^

Pol α^Cat^ contains the DNA polymerase active site mutations Pol 1–D996S, D998S and was purified as previously described in (Georgescu et al., 2015) with few modifications. Briefly, we inserted a 2× strep tag at the N-terminus of Pol 12 for elimination of N-terminally proteolyzed Pol 12. After batch purification from anti-FLAG agarose (Sigma) the eluent was mixed with 300 μL of StrepTactin resin (IBA Lifesciences) equilibrated with PBS buffer supplemented with 10% (*v/v*) glycerol and washed. Pol α^Cat^ was eluted in PBS buffer supplemented with 10% (*v/v*) glycerol and 10 mM desthiobiotin. Fractions were pooled, frozen in liquid N_2_ and stored in aliquots at −80°C.

##### Production of SFP synthase reagents for protein labeling

SFP synthase was purified by a nickel-NTA chromatography as previously described (Yin et al., 2006). Alexa Fluor 488 (AF488) was functionalized by Co-enzyme A (CoA) and purified by HPLC as previously described (Yin et al., 2006) and an LD650–CoA (a CoA-derivatized photostable version of Cy5) was purchased by Lumidyne Technologies (USA).

##### Preparation of LD650–Pol ε and AF488–Pol ε

To obtain Pol ε labeled with a single LD650 (Lumidyne Technologies) or Alexa Fluor 488 (Invitrogen) dye, we inserted the “S6” peptide (GDSLSWLLELLN) (Zhou et al., 2007) between the N-terminus of Pol 2 and its 3× FLAG tag. The resultant Pol ε–S6 plasmid was overexpressed and purified in *S. cerevisiae* as previously described (Georgescu et al., 2014). For labeling Pol ε–S6, SFP enzyme, and either LD650–CoA or AF488–CoA were then incubated at a 1:2:5 molar ratio for 1 hour at room temperature in the presence of 10 mM MgCl_2_. Excess dye and SFP enzyme were removed by purification on a Superose 6 column (GE Healthcare) with a buffer containing 25 mM Tris-HCl pH 7.5, 150 mM NaCl, 1 mM dithiothreitol, 5% (*v/v*) glycerol. Fractions were frozen in liquid N_2_ and stored in aliquots at −80°C. The degree of labeling was determined to be ~100% for both LD650–Pol ε and A488–Pol ε by UV/vis spectrophotometry.

##### Preparation of LD650–CMG

CMG labeled with LD650 (Lumindyne Technologies) was produced as previously described in (Wasserman et al., 2019).

##### Preparation of Cy5–Pol δ

Pol δ was prepared in *S. cerevisiae* as previously described (Georgescu et al., 2014) and subsequently dialyzed into 250 mM NaCl, 5% (*v/v*) glycerol, 50 mM potassium glutamate, 25 mM HEPES pH 7.5, and 2 mM dithiothreitol. Following dialysis Pol δ was labeled with a 5-fold molar excess of Cy5–NHS (GE Healthcare) for 5 min at 4°C. Excess dye was removed by five buffer exchange steps through a centrifugal filter (50K MWCO; Millipore) following the manufacturer’s instructions. Labeled Cy5–Pol δ was frozen in liquid N_2_ and stored in aliquots at −80°C. The degree of labeling was measured to be 5 fluorophores per Pol δ holoenzyme by UV/vis spectrophotometry.

##### Preparation of AF647–Pol δ^32-^

Alexa Fluor 647 (Invitrogen), was used to label Pol δ^32-^. Labeling reactions were carried out using 3-fold molar excess of dye with 58.5 μM Pol δ^32-^ in 320 μL of Pol δ^32-^ labeling buffer (30 mM Tris-HCl pH 7.6, 2 mM dithiothreitol, 300 mM NaCl, 50 mM potassium glutamate, 10% (*v*/*v*) glycerol) for 10 min at 4°C with gentle rotation. Immediately following the coupling reaction, excess dye was removed by sequential elutions from two 0.5 mL Zeba spin desalting columns (7K MWCO; Thermofisher) following the manufacturer’s instructions equilibrated in buffer containing 30 mM Tris-HCl pH 7.6, 2 mM dithiothreitol, 300 mM NaCl, 50 mM potassium glutamate, 10% (*v*/*v*) glycerol. Labeled AF647–Pol δ^32-^ was frozen in liquid N_2_ and stored in aliquots at −80°C. The degree of labeling was measured to be 1 fluorophore per Pol δ^32-^ holoenzyme by UV/vis spectrophotometry.

##### Preparation of AF647–RPA

Alexa Fluor 647 (Invitrogen) was used to label RPA. Labeling reactions were carried out using 5-fold molar excess of dye with 45 μM RPA in 550 μL of RPA labeling buffer (50 mM Tris-HCl pH 7.6, 3 mM dithiothreitol, 1mM EDTA, 200 mM NaCl, 10% (*v*/*v*) glycerol) for 2 hours at 23°C with gentle rotation. Immediately following the coupling, excess dye was removed by gel filtration at 1 mL/min through a column (1.5 × 10 cm) of Sephadex G-25 (GE Healthcare), equilibrated in gel filtration buffer (50 mM Tris-HCl pH 7.6, 3 mM dithiothreitol, 1mM EDTA, 200 mM NaCl, 20% (*v*/*v*) glycerol). Labeled AF647–RPA was frozen in liquid N_2_ and stored as single use aliquots at –80°C. The degree of labeling was measured to be 1 fluorophore per RPA trimer by UV/vis spectrophotometry.

#### DNA Substrate Preparation

##### Oligonucleotides and DNA

Oligonucleotides were purchased from Integrated DNA technologies (USA). Plasmid pSuperCos1 DNA was purified by Aldevron (USA).

##### Linear forked doubly-tethered DNA substrates

To make the doubly tethered linear fork DNA substrate, plasmid pSupercos1 DNA (van Loenhout et al., 2012) was linearized overnight at 37°C with 100 U of *Bst*XI in 1 x Buffer 3.1 (New England Biolabs). The 18,284 bp fragment was purified with a Wizard SV gel and PCR clean up kit (Promega) and the concentration was measured. The fork junction was constructed by annealing 15.3 pmol of 160Ld, 91.8 pmol 99Lg, 1530 pmol of fork primer (Table S1) by heating at 94°C for 5 min before slowly cooling. Similarly, the biotinylated blocking duplex was generated by annealing 5.3 pmol of blockingLd and blockingLg by heating at 94°C for 5 min before slowly cooling. 1.5 pmol of the 18,284 bp linear DNA template was ligated to the pre-annealed fork junction and biotinylated blocking duplex in 1 x T4 ligase buffer (New England Biolabs) and 2000 U of T4 ligase (New England Biolabs) overnight at 16°C. The ligated linear forked DNA substrates were purified from excess DNA oligonucleotides by adjusting NaCl to 300 mM and loaded by gravity onto a Sepharose 4B (Sigma; 1 × 25 cm) column, equilibrated in gel filtration buffer (10 mM Tris-HCl pH 8.0, 1 mM EDTA, and 300 mM NaCl). Ligated biotinylated linear DNA substrates eluted as a single peak in the column void volume, fractions under the peak were analysed by agarose gel electrophoresis. Fractions were pooled and dialysed overnight in 2 L of sterilized TE buffer, concentrated 2-fold in a vacuum concentrator and the concentration measured. Aliquots were stored at –80°C.

##### Primed linear DNA substrate

The primed linear substrate used to test the polymerase activity of Pol α^Cat^ (Fig S7A) was constructed as follows (Table S1). To create the 243 nt template, Near 143mer and Far 100mer were ligated together by first mixing with Near Far bridge in a 1:1:3 molar ratio in the presence of 50 mM NaCl and 5 mM Trisodium Citrate pH 7.0. Next, the duplex was annealed by heating to 94°C for 5 min before slowly cooling. Following this 1 mM ATP and 4,000 Units of T4 Ligase (New England Biolabs) were added and the reaction was incubated for 16 hours at 15°C. The ligated product was purified on an 8% polyacrylamide, 8M Urea denaturing gel using SYBR Safe stain (Invitrogen) to visualize the single-stranded product. The DNA was recovered by crushing the gel and soaking in buffer TE pH 8.0 for 16 hours at room temperature. Followed by spinning at 15,000 rpm for 5 minutes, to recover the supernatant. 30-mer Primer was 5’-end labeled with γ-^32^P-ATP by T4 PNK (New England Biolabs) according to manufacturer instructions and purified on an S-200 HR microspin column (GE Healthcare). The radiolabeled primer was annealed to the 243 nt template in a 2:3 ratio.

#### Ensemble DNA replication assay

To confirm the absence of DNA polymerase activity of Pol α^Cat^ (Fig S7B), 2 nM of the linear DNA substrate primed with a 5’-^32^P-primer (see Primed linear substrate) was incubated with either Pol α or Pol α^Cat^ in the presence of 25 mM Tris-OAc pH 7.5, 5% (*v/v*) glycerol, 100 μg/mL BSA, 5 mM TCEP, 10 mM Magnesium Acetate, 50 mM Potassium glutamate, and 0.1 mM EDTA. Replication reactions were initiated by the addition of 80 μM each of dTTP, dATP, dCTP, and dGTP and allowed to proceed for 20 minutes at 30°C. Reaction volumes were 20 μL. Reactions were quenched by mixing with an equal volume of 2× stop buffer containing 80% (*w/v*) *formamide*, 8 mM EDTA, and 1% (*w/v*) SDS and were subsequently run on a 10% polyacrylamide, 8M Urea denaturing gel for 1.5 hours at 125 V. The gel was exposed to a phosphorimaging screen for 16 hours and visualized with a Typhoon FLA 9500 scanner (GE Healthcare).

#### Single-molecule DNA replication assays

##### Flow cell preparation

Flow cells were prepared as described previously (Geertsema et al., 2015; Lewis et al., 2017a). Briefly, a polydimethylsiloxane (Sylgard) lid was placed on top of a PEG-biotin-functionalized microscope slide (24 × 24 mm, Marienfeld) to create a 1-mm-wide and 100-μm-high flow channel (volume 1 μL). Polyethylene tubes (PE-60: 0.76-mm inlet diameter and 1.22-mm outer diameter, Walker Scientific) were inserted to allow for a buffer flow. To help prevent nonspecific interactions of proteins and DNA with the surface, the chamber was blocked with blocking buffer (50 mM Tris-HCl pH 7.6, 50 mM KCl, 2% (*v/v*) Tween-20). The forked DNA substrates (20 pM) were flowed through the chamber for 20 min at 17 μL/min in the presence of 200 μM Chloroquine (Sigma). The DNA was visualized by flowing in replication buffer (25 mM Tris-HCl, pH 7.6, 10 mM magnesium acetate, 50 mM potassium glutamate, 40 μg/mL BSA, 0.1 mM EDTA, 5 mM dithiothreitol, and 0.0025% (*v/v*) Tween-20) with 150 nM S.O. (Life Technologies).

##### Replication reaction conditions

Conditions for the pre-assembly replication reactions were carried out in three steps. First, 30 nM CMG was loaded at 10 μL/min in CMG loading buffer with 60 nM Mcm10 and 400 μM ATP. Following this, the replisome was assembled by introducing 20 nM Pol ε, 20 nM Pol δ, 20 nM Pol α, 20 nM Ctf4, 20 nM PCNA, 20 nM RFC, and 30 nM MTC in replication buffer supplemented with 400 μM ATP, and 60 μM dCTP/dGTP at 10 μL/min for 5 min. Replication was initiated by washing the flow cell with 100 μL (100 flow cell volumes) at 50 μL/min with a solution containing 60 nM Mcm10, 20 nM PCNA, 20 nM RFC, 200 nM RPA, and 30 nM MTC in replication buffer supplemented with 5 mM ATP, 125 μM dCTP, dGTP, dATP, and dTTP, and 250 μM CTP, GTP, ATP, and UTP, and 150 nM S.O.

Conditions for replication under the continuous presence of all proteins were performed in multiple stages. First, 30 nM CMG (or LD650–CMG) was loaded at 10 μL/min in CMG loading buffer (25 mM Tris-HCl, pH 7.6, 10 mM magnesium acetate, 250 mM potassium glutamate, 40 μg/mL BSA, 0.1 mM EDTA, 5 mM dithiothreitol, and 0.0025% (*v/v*) Tween-20), with 60 nM Mcm10 and 400 μM ATP. When LD650–CMG was used, the flowcell was subsequently washed under a continuous flow of CMG loading buffer supplemented with 500 mM NaCl at 10 μL/min for 10 min. Then, replication reactions were initiated by introducing 60 nM Mcm10, 20 nM Pol ε (unless specified otherwise), 20 nM Pol δ (unless specified otherwise), 20 nM Pol α (unless specified otherwise), 20 nM Ctf4, 20 nM PCNA, 20 nM or 2nM RFC, 200 nM RPA, and 30 nM MTC in replication buffer supplemented with 5 mM ATP, 125 μM dCTP, dGTP, dATP, and dTTP, and 250 μM CTP, GTP, ATP, and UTP, and 150 nM S.O. In CMG challenge experiments 10 nM CMG was added to the replication reaction.

##### Imaging conditions

All single-molecule assays were carried out on an inverted microscope (Nikon Eclipse Ti-E) fitted with a CFI Apo TIRF 100× oil-immersion objective (NA 1.49, Nikon). The temperature was maintained at 31.2°C by an electrically heated chamber (Okolab). dsDNA was visualized every 10 s for 30 min by exciting the S.O. with a 568-nm laser (Coherent, Sapphire 568–200 CW) at 80 mW/cm^2^. The red fluorescently labeled proteins were excited at 80 mW/cm^2^ (800 W/cm^2^ during a FRAP pulse) with a 647-nm laser (Coherent, Obis 647– 100 CW). The AF488–Pol ε was visualized with a 488-nm laser at 140 mW/cm^2^. The signals were spectrally separated using appropriate filter sets (Chroma) and fluorescence signals collected on an Evolve 512 Delta EMCCD (Photometics). Typically, nine fields of view (five for the FRAP experiments) were selected for imaging. Single-molecule experimental results were derived from at least three or four technical replicates for each experimental condition.

#### Determination of RPA binding footprint

Flow cells were prepared as described above. Oligonucleotides 57-mer or 99Lg (Table S1) were incubated with AF647–RPA in replication buffer for 10 min at 25°C. DNA–RPA complexes were introduced on the surface of the flow cell and washed with 100 μL of replication buffer. The AF647–RPA were excited at 80 mW/cm^2^ with a 647-nm laser (Coherent, Obis 647–100 CW). Imaging was carried out as described in ‘Single-molecule DNA replication assays’.

#### Measurement of the stoichiometry of fluorescently labeled proteins at the replisome

The average intensity of labeled proteins (Pol ε, Pol δ, CMG, or RPA) was quantified by immobilization on the surface of a cleaned microscope coverslip in replication buffer at 6 pM. Imaging was carried out under the same conditions used during the single-molecule replication experiments. We calculated the integrated intensity for every fluorescent protein in a field of view after applying a local background subtraction (Lewis et al., 2017b). The histograms obtained were fit with a Gaussian distribution function using MATLAB 2016b, to give a mean intensity. We calculated the total number of molecules at every time point during DNA replication by dividing their intensities by the intensity of a single molecule. Subsequent histograms were fit to Gaussian distribution using MATLAB 2016b.

#### Analysis of single-molecule replication kinetics and protein dynamics

All analyses were carried out using ImageJ/Fiji (1.51w) and Matlab 2016b, and in-house built plugins. The rate of replication of a single molecule was obtained by first tracking the position of the leading-strand spot using the Linear-motion LAP tracker in TrackMate v3.6.0 (Tinevez et al., 2017). Individual rate segments were identified using kinetic change-point analysis (Duderstadt et al., 2016; Hill et al., 2018; Lewis et al., 2017b). The rates obtained from this algorithm were weighted by the DNA segment length, to reflect the number of nucleotides that were synthesized at this rate. This places more significance on the longer rate segments, as they have a higher signal-to-noise ratio compared with shorter segments (Lewis et al., 2017b).

To measure the intensity of the leading-strand spot at the replication fork, we tracked the position of the leading-strand spot and integrated the intensity for all colours simultaneously over time. To obtain the characteristic exchange time τ from the FRAP experiments, the data were fit with a FRAP recovery function correcting for photobleaching (Beattie et al., 2017; Lewis et al., 2017a; Spenkelink et al., 2019) (Formula 1, where *a* is the amplitude of photobleaching, t_b_ is the photobleaching time (measured in Figure S6), and I_0_ is the number of polymerases at the fork at steady state).

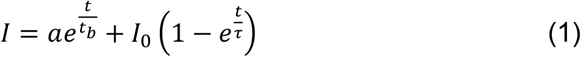

The maximum exchange rate was obtained by fitting the data with a hyperbolic equation (Formula 2, where R is the exchange rate, R_max_ is the maximum exchange rate, [Pol] is the polymerase concentration, and K_b_ is the characteristic binding constant).

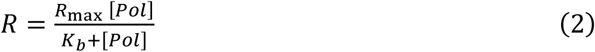

### QUANTIFICATION AND STATISTICAL ANALYSIS

The number of molecules or events analyzed is indicated in the text or figure legends. Errors reported in this study represent the standard error of the mean (S.E.M) or the error of the fit, as indicated in the text or figure legends. Every single-molecule replication experiment was carried out at least in triplicate.

### DATA AND SOFTWARE AVAILABILITY

Raw data is available upon request. All home-built ImageJ plugins used in this study are freely available on the Github repository for Single-molecule/Image analysis tools (https://github.com/SingleMolecule) or available upon request.

## Supplemental Information

**Table S1.**
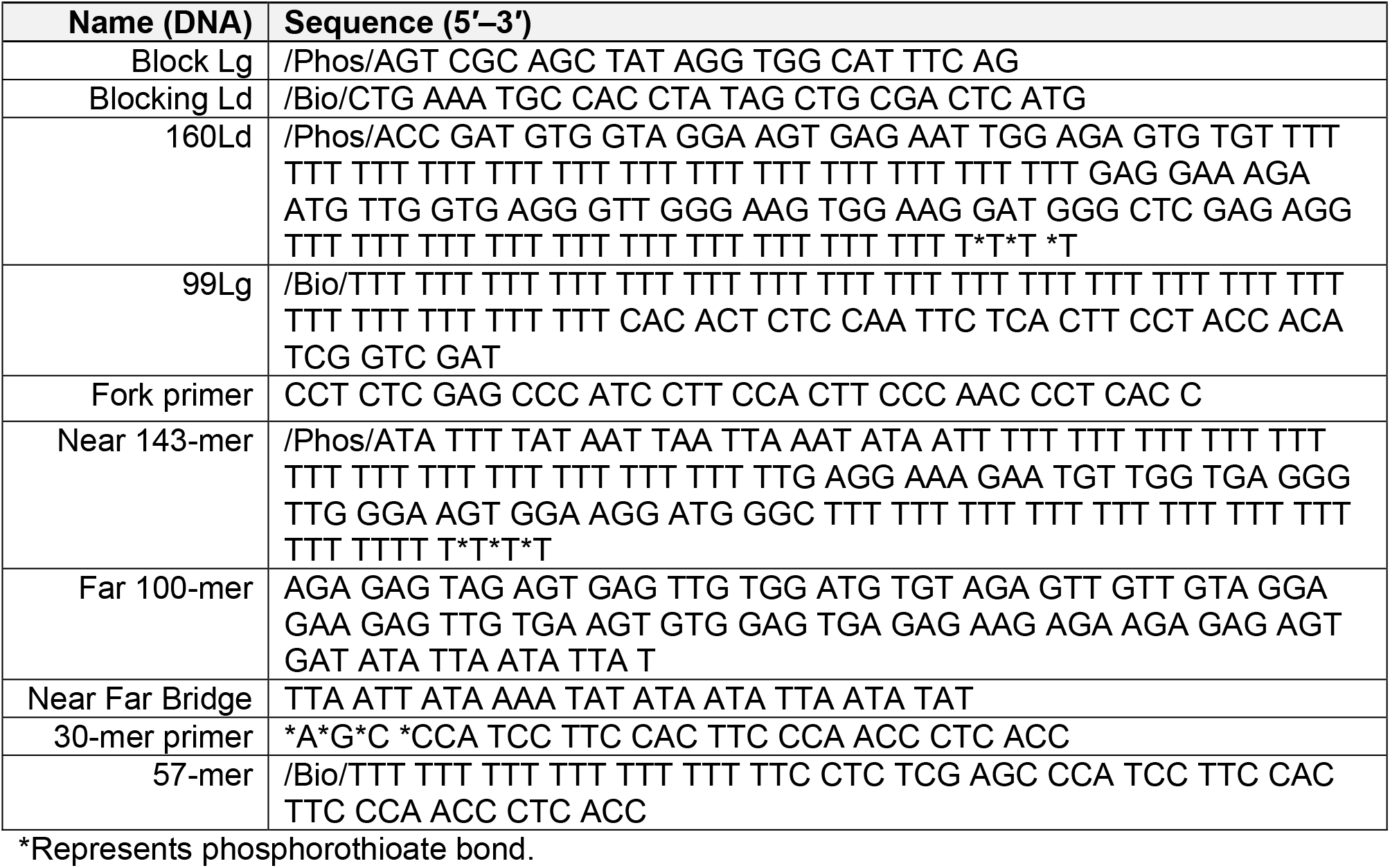
Sequences of oligonucleotides used in this study.

**Figure S1.**
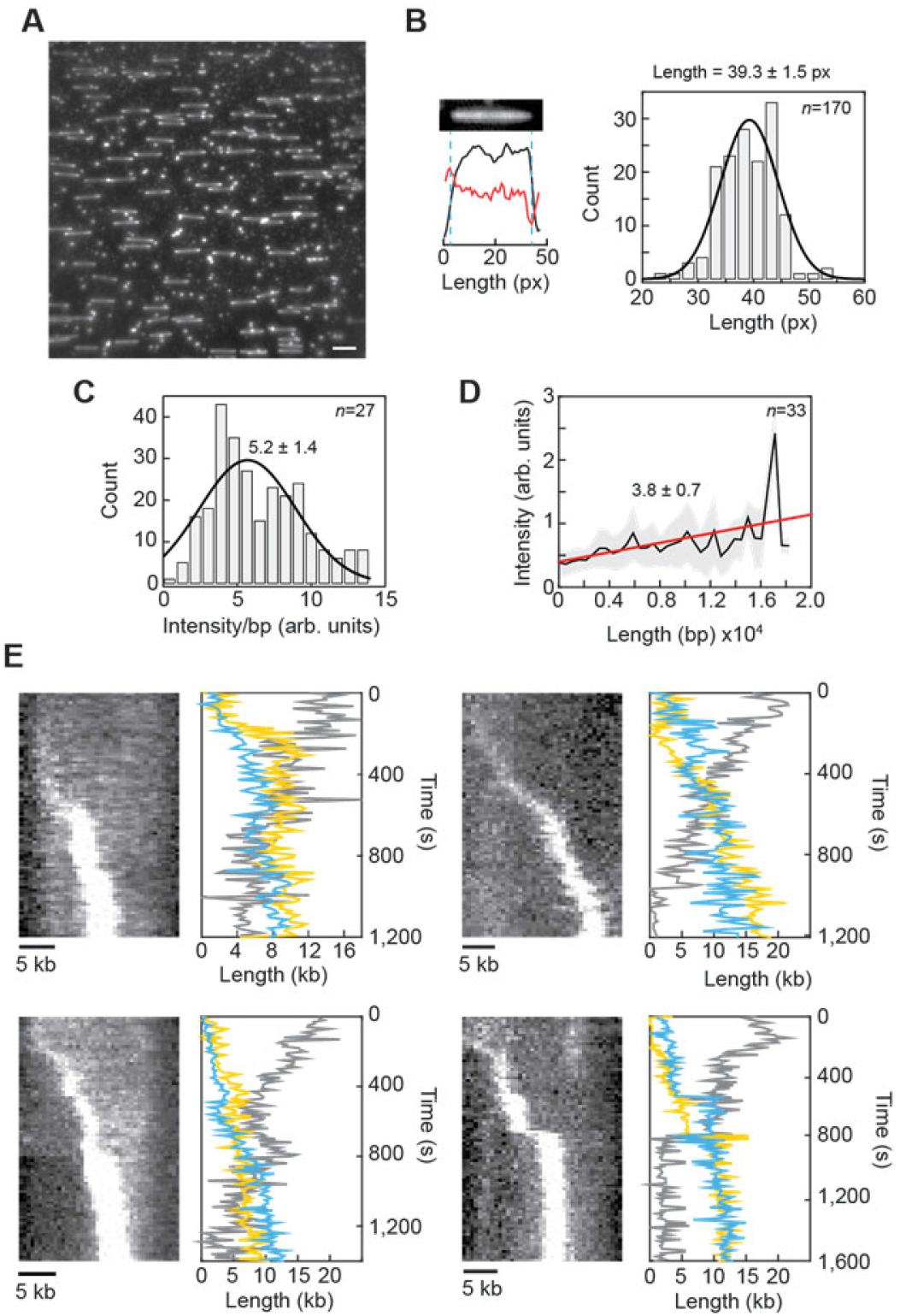
Characterization of DNA replication on linear dsDNA substrates, Related to Figure 1. (A) Representative field of view showing doubly tethered DNA substrates in the absence of any buffer flow. Scale bar = 20 kb. (B) Calibration of the length of the DNA substrates under our experimental conditions. Error represents S.E.M. *n* = number of molecules. (C) Fluorescence intensity per base pair in the leading-strand spot. (D) Fluorescence intensity per base pair from the 18.3 kb substrate. Error represents S.E.M. *n* = number of molecules. (E) Representative kymographs and corresponding graphs showing length of the lagging-strand (yellow), leading-strand (blue) and parental DNA (gray) as a function of time, measured by the integrated intensity of the dsDNA.

**Figure S2.**
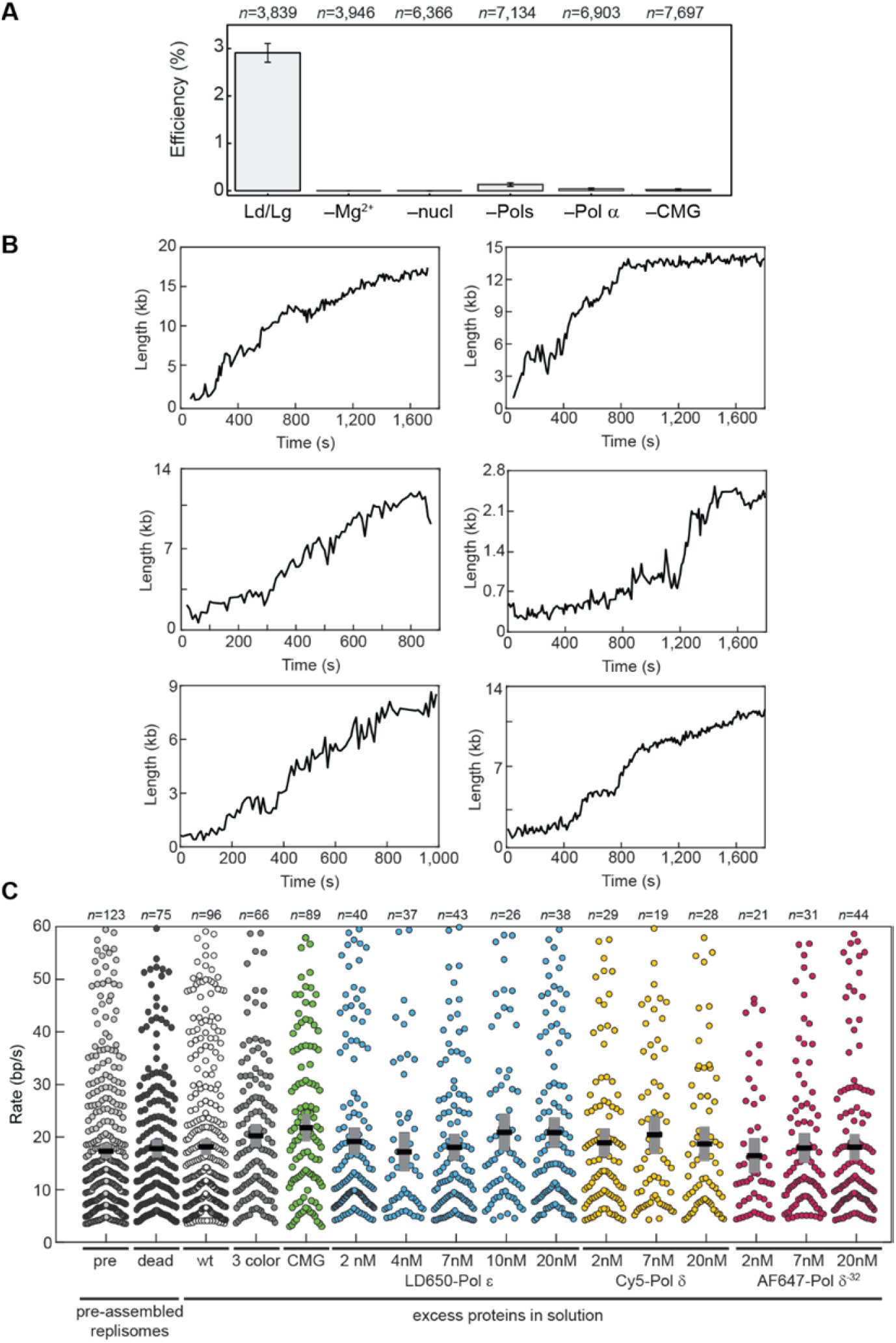
Validation and comparison of reaction conditions, Related to Figures 1–5. (A) Negative controls. The replication efficiency was defined as the number of templates that showed replication, divided by the total number of observed DNA substrates. Error bars represent S.E.M. *n* = number of DNA substrates. (B) Example trajectories. Distance travelled as a function of time for six representative single molecules. (C) Comparison of replication rates at different concentrations of polymerases. Scatter plots of the single-molecule rate segments from pre-assembled replisomes (gray) and Pol α^Cat^ (black); Replisomes challenged with excess proteins in solution, using wild-type polymerases (white), 20 nM AF488–Pol ε and Cy5–Pol δ (dark gray), LD650–CMG (green) increasing concentrations of LD650–Pol ε (blue), increasing concentrations of Cy5–Pol δ (yellow), and increasing concentrations of AF647–Pol δ^32-^ (red). The black line and gray box represent the mean and S.E.M. *n* = the number of molecules.

**Figure S3.**
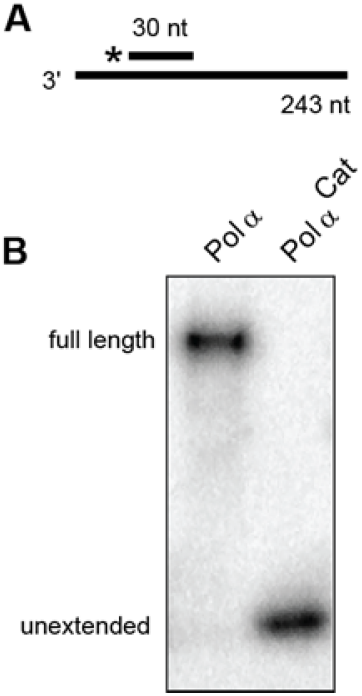
Pol α^Cat^ is unable to carry out DNA polymerase activity, Related to Figure 2. (A) Schematic representation of the 243 nt primed linear DNA substrate. (B) Ensemble DNA replication assay (Lane 1) wild-type Pol α replicates the substrate to full length. (Lane 2) Pol α^Cat^ containing mutations in Pol 1–D996S,D998S is unable to extend the primer.

**Figure S4.**
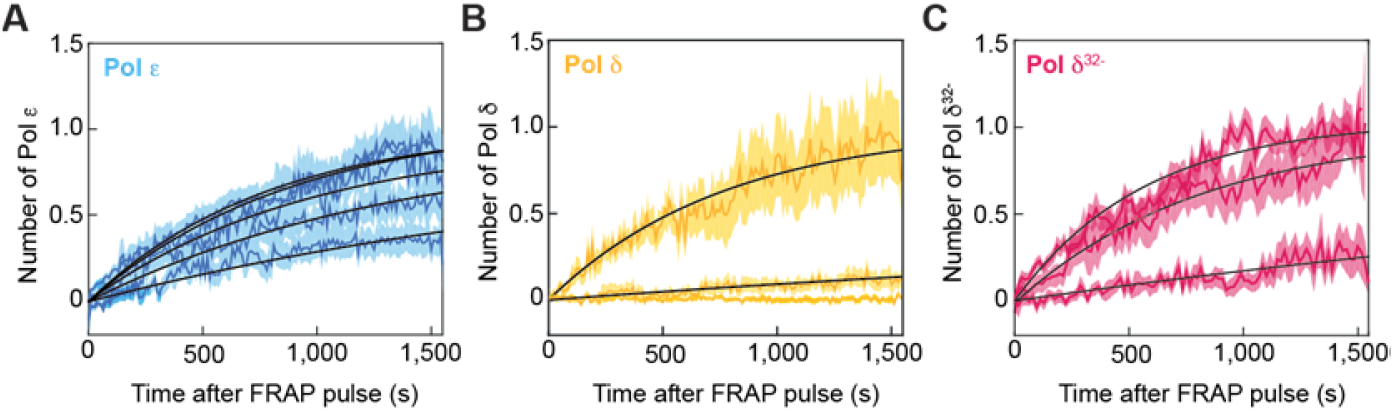
FRAP recovery curves for labeled polymerases, Related to Figures 3 and 5. (A) FRAP recovery curves for LD650–Pol ε. Average number of LD650–Pol ε at the replication fork over time after the FRAP pulse for 20, 10, 7, 4, and 2 nM of LD650–Pol ε. (B) FRAP recovery curves for Cy5–Pol δ. Average number of Cy5–Pol δ at the replication fork over time after the FRAP pulse for 20, 7, and 2 nM of Cy5–Pol δ. (C) FRAP recovery curves for AF647–Pol δ^32-^. Average number of AF647–Pol δ^32-^ at the replication fork over time after the FRAP pulse for 20, 7, and 2 nM of AF647–Pol δ^32-^. All recovery curves are fit using formula 1 (See methods for detailed explanations). Error represents Standard deviation.

**Figure S5.**
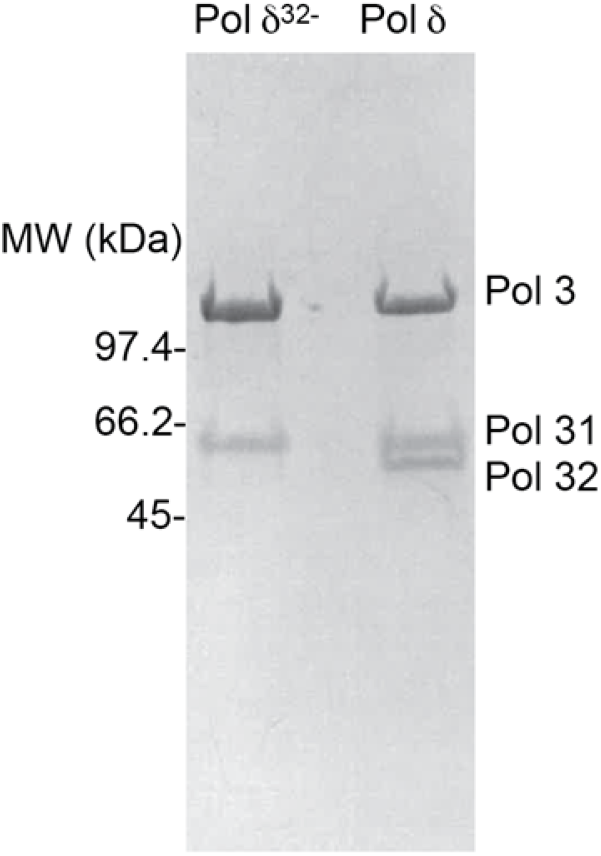
Coomassie blue stained SDS-PAGE gel of purified Pol δ and Pol δ^32-^, Related to STAR Methods. (Lane 1) Pol δ holoenzyme lacking subunit 32 and (Lane 2) Pol δ holoenzyme.

**Figure S6.**
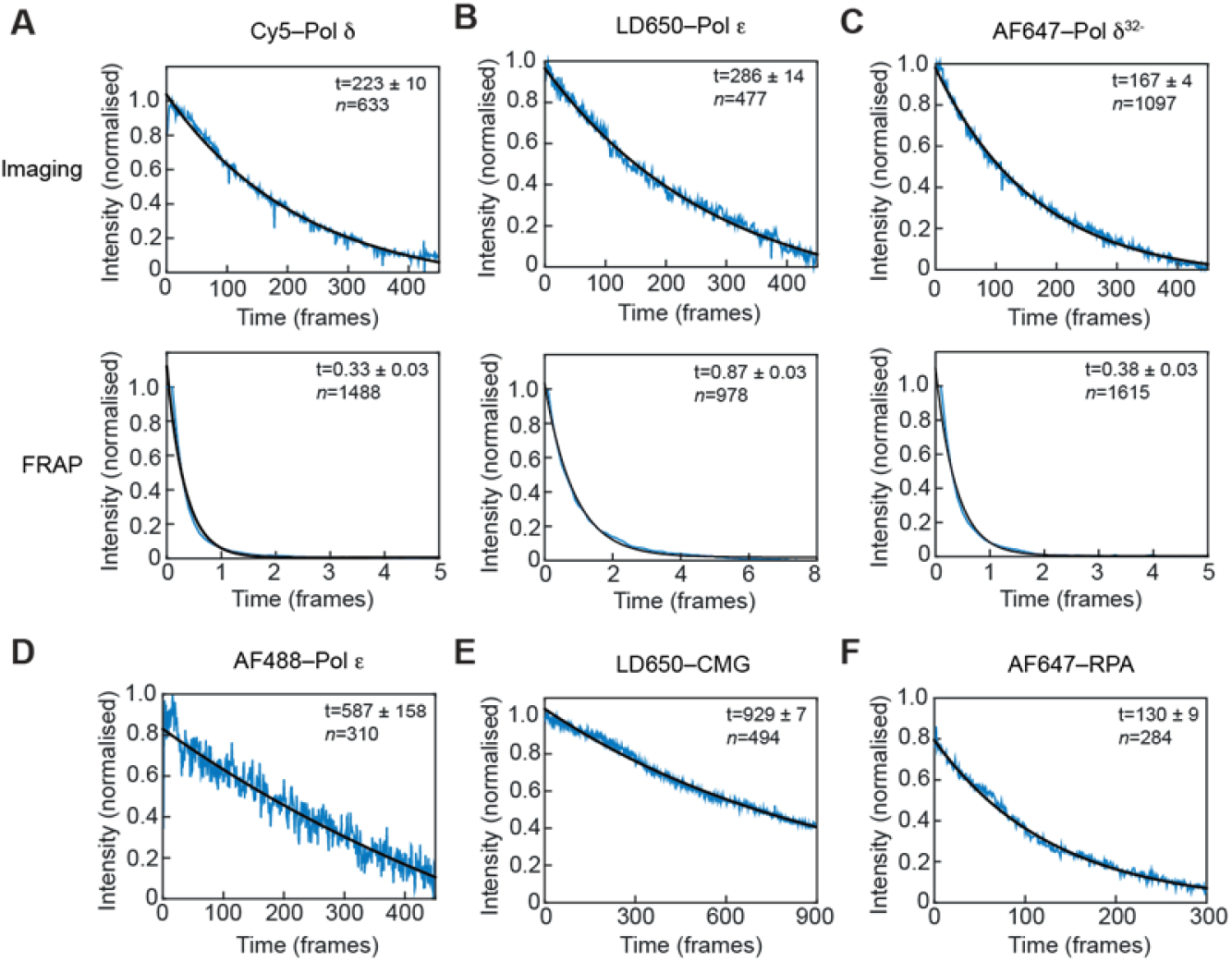
Quantification of photobleaching kinetics of fluorescently labeled proteins, Related to STAR Methods. The normalized average intensity as a function of time (in frames) under imaging conditions (top) and FRAP photobleaching-pulse conditions (bottom) for (A) Cy5–Pol δ (B) LD650–Pol ε, (C) AF647–Pol δ^32-^(D) The normalized average intensity as a function of time (in frames) under imaging conditions for AF488–Pol ε, and (E) LD650–CMG (F) AF647–RPA. Errors represent error of the fit.

**Movie S1 Single-molecule movie showing DNA replication on a single DNA substrate from example kymograph in Figure 1B**. Scale bar = 1 μm.

